# A Computational Framework for Pulmonary Assessing Wave Intensity Following Simulated Lung Resection

**DOI:** 10.64898/2026.03.16.712097

**Authors:** Jay A. Mackenzie, Nicholas A. Hill

**Affiliations:** School of Mathematics & Statistics, University of Glasgow, Glasgow, G12 8QQ, UK

## Abstract

**Background and Objectives:** Lung cancer is one of the most frequently diagnosed cancers worldwide. While non-surgical treatment options have increased in number and efficacy, lung resection for primary cancers is still a mainstay of treatment. Lung resection has been shown to impair right ventricular function, although the mechanism for the impairment remains unclear. Wave intensity is increasingly used as a metric for increased post-operative afterload. Here, we develop a computational framework to assess the impact of simulated lung resection on wave intensity to establish that post-operative changes in wave intensity are attributable to the change in pulmonary artery morphometry.

**Methods:** We analyse a 48 pulmonary arterial surfaces segmented from CT images in patients with no evidence of lung disease to obtain 1D representations of the pulmonary vasculature. For each pulmonary vasculature we sequentially remove vessel branches to mimic post-operative morphometric changes to the arterial network. Using an established 1D computational flow model, we simulate pulsate blood flow in 44 pre-operative cases and 1596 post-operative cases. We compute wave intensity in the main, right, and left pulmonary arteries for all simulations.

**Results:** We compare the change in computed wave intensities pre-versus post-operatively to the results of an experimental clinical study comparing pre- and post-operative wave intensity in a 27 patient cohort. We see good agreement between the changes in the parameters of wave intensity between this study and those reported in the clinical study. Further, we capture flow distribution the changes pre-versus post-operatively which indicates that the computational model behaves as expected.

**Conclusions:** In this preliminary study on a computational framework to capture changes in pulmonary arterial haemodynamics following lung resection, we have shown that our model and analysis pipeline is capable of capturing post-operative changes to wave intensity and flow redistribution between the pulmonary arteries following lung resection. These results motivate further research to develop and validate a patient specific model which is an area of active research for us.

## 1. Introduction

In 2022, lung cancer was the most frequently diagnosed cancer worldwide [1] with 2.48 million new cases and 1.8 million deaths, respectively. While incidence of lung cancer varies regionally [2], it is the most commonly diagnosed cancer in Scotland[3]. While non-surgical treatment options have increased in number and efficacy [4], lung resection to treat or manage disease remains relatively common, but regionally variable, with rates between 13% and 32% for all lung cancers [5].

Lung resection has been shown to negatively impact right ventricular function [6]. According to Glass et al. [7] This impairment is typically attributed to increased afterload following resection but previous studies have failed to show increased pulmonary artery (PA) pressure or vascular resistance post-operatively. They suggest analysis of PA wave reflections via wave intensity analysis (WIA) to assess the contribution to pulsatile after-load. Glass et al. [6] applied wave intensity analysis to the MRI derived flow and area measurements from 27 lung resection patients pre-operatively and on post-op day 2. They find a marked changes to wave intensity in the pre-versus post-operative measurements in the operative and non-operative PAs. These changes are summarised in Table 3. It is currently unclear whether these changes occur due to some biological compensatory mechanism or due to the change in the geometry of the pulmonary arterial tree.

In this hemodynamic modelling study, we develop a novel computational pipeline to simulate flow in 44 pre-operative PA geometries obtained from computed tomography (CT) images. We make rational and systematic changes to these PA networks to obtain 1639 simulated post-operative PA trees. We simulate blood flow in all 1683 networks and perform the same processing steps and WIA as Glass et al. [8, 6]. We compare the results of these simulation and processing steps against the measured data presented by Glass [8] and conclude that the computational model behaves similarly to the measured flows. This is indicative that post-operative changes in PA wave intensity are due to the alteration in PA geometry caused by the resection.

## 2. Methods

The hemodynamic model comprises a few parts: a blood vessel network, a set of model equations, and a set of boundary conditions. We address each of these in turn.

In this study, we employ the mathematical and computational model developed by Vaughan [9], extended from Olufsen [10, 11], to study pulse propagation in the pulmonary arterial tree before and after simulation lung resection. Lung resection is simulated by removing vessels from the large arterial tree. In the model, large vessels and their branching are described explicitly, and small vessels are described implicitly. If a vessel is not large then it is small. Large vessels typically make up the first few generations of the bifurcating arterial tree after which self-similar diverging trees of small vessels are generated. These self-similar vessel networks are known as structured trees, and can be thought of as an average description of a vascular bed [10]. They are truncated at a specified minimum radius and they have specified properties such as wall stiffness, length-to-radius ratio, and branching asymmetry.

The large vessel networks used in this study come from a pre-segmented data set of chest CTs in patients with no evidence of lung disease [12]. The pulmonary veins were not segmented, so are not included in this study as in Olufsen et al. [11]. Fuller details are given in Appendix A.1.

### 2.1. Boundary Conditions

As mentioned above, much of the vasculature is captured using a structured tree approach. This is the down-stream boundary condition on the model. The numerous vessels in the structured trees provide resistance to upstream flows. This boundary condition and parameters used to implement it are given in Appendix A.2 for completeness. The upstream boundary condition given as the flow (mLs^−1^) over a single cardiac cycle. The period of the cardiac cycle is 0.7 s. In this study, we use the inflow profile presented by Olufsen et al. [13] which is imposed at the inlet of the main pulmonary artery (MPA) in all cases. We choose to use a flow based boundary condition, as opposed to a pressure based boundary condition as total blood volume will revert to a baseline post-surgery. Further, we are interested in simulating increased right ventricular load in the form of increased pressure in the MPA, and fixing pressure as a boundary condition would obfuscate such findings. The inlet flow profile used in this study can be seen in Fig. 1.

**Figure 1:**
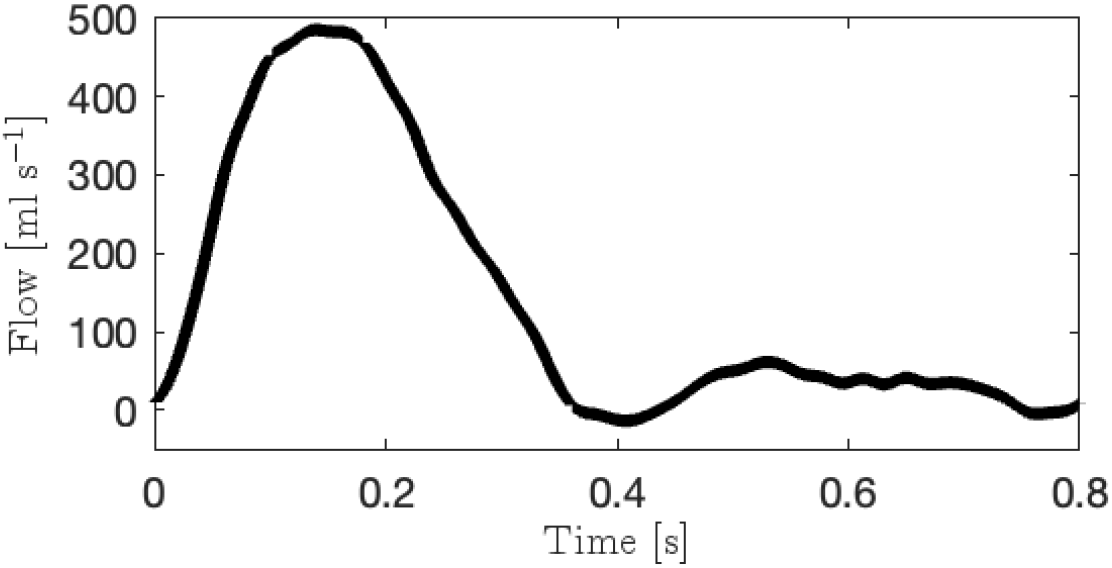
Inflow profile at the inlet of MPA. The profile is interpolated from MRI measurements sampled at 45 points per period and averaged over 5 cardiac cycles. From Olufsen et al. [13].

### 2.2. Computational Domain

In this section, we discuss the process of obtaining 1D computational domains from segmented pulmonary CT images.

#### 2.2.1. Data

The large arterial networks used here are derived from imaged data. These data were collected and segmented as per Goubergrits et al. [12] who have made available the segmented pulmonary surfaces for 48 aortic stenosis patients. The data were made available in `stl` format. The silhouettes of these surfaces can be seen in Figs. 2 and 3 as projections onto the coronal plane. These data are a collection of 3D surfaces, and the computational model is a 1D scheme, so we essentially need to transform the surfaces into a collection of frusta of a cone in which each frusta best represents a vessel branch. Here, we discuss this transformation from surface to 1D computational domain.

**Figure 2:**
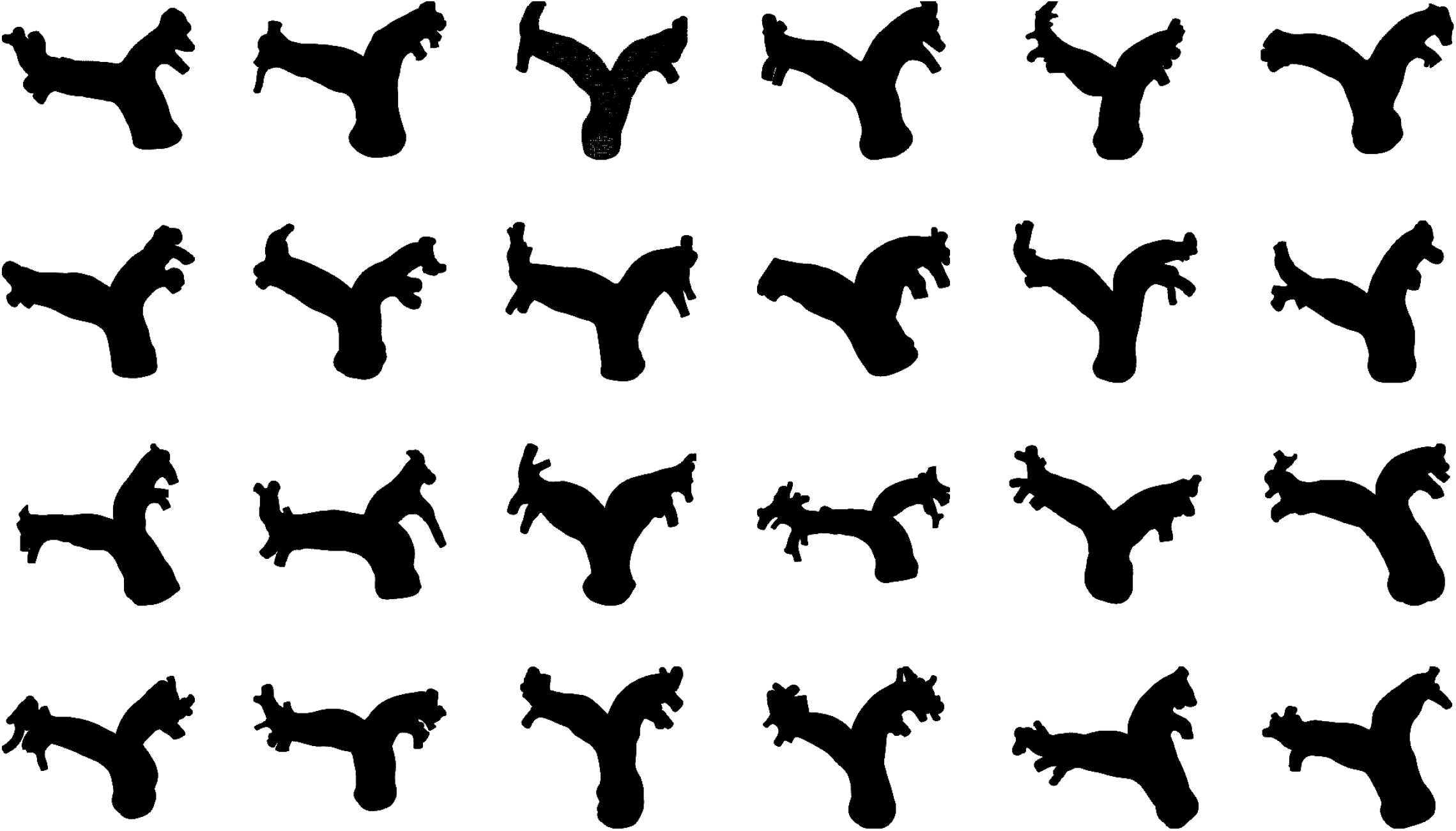
Coronal projections of 24 of the human pulmonary arterial trees collected and analysed by Goubergrits et al. [12].

**Figure 3:**
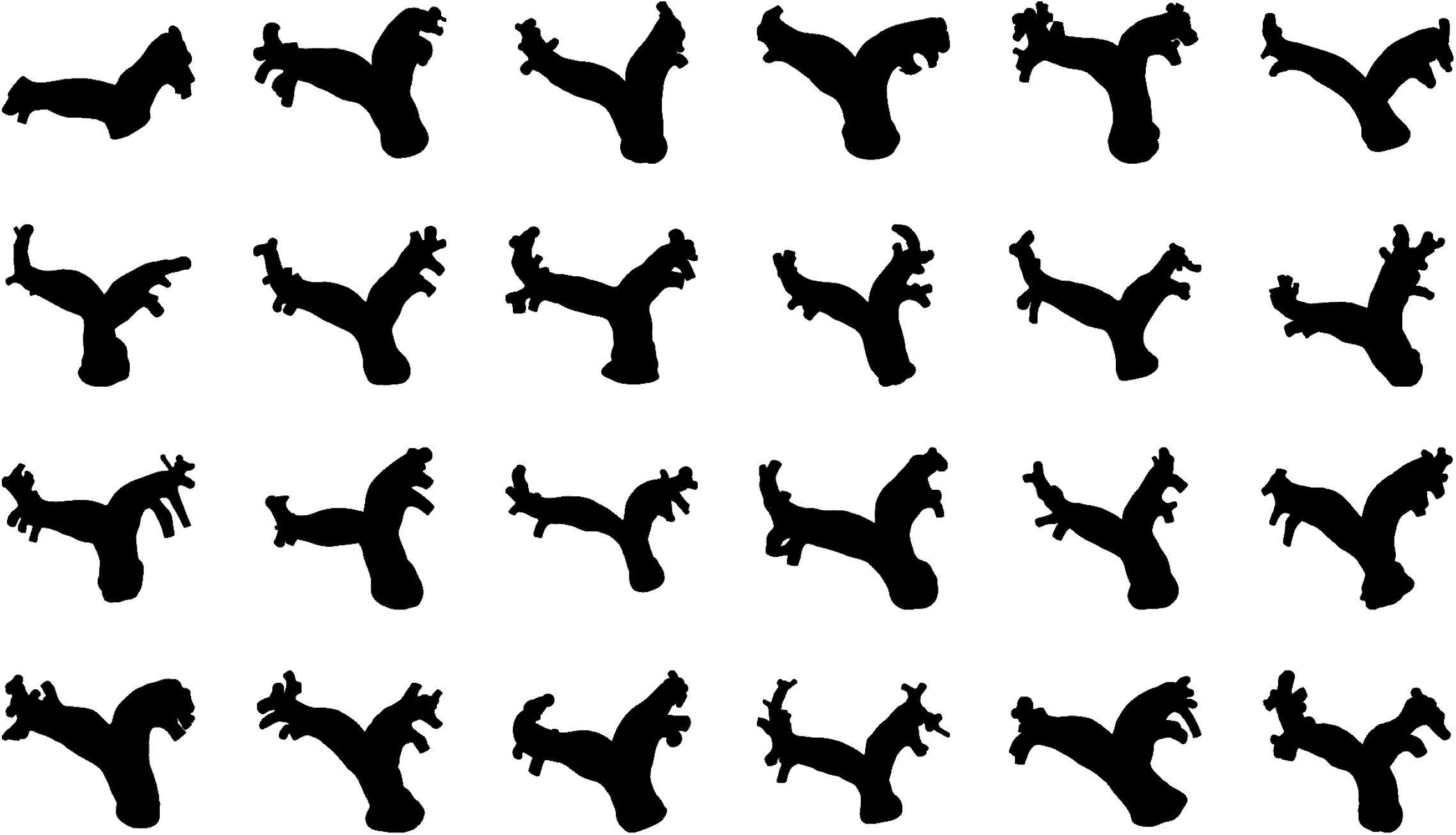
Coronal projections of 24 of the human pulmonary arterial trees collected and analysed by Goubergrits et al. [12].

Initially, we use the vascular modelling tool kit (VMTK)[14] to find the centrelines for each tree that run from the inlet of the MPA to the terminii of each of the outlet vessels. The inlet and outlet surfaces are detected automatically by VMTK. The inlet vessel is manually specified by the user to ensure that all centrelines begin at the proximal end of the MPA. The correct choice was always obvious because of vessel calibre and branching structure. The centrelines are converted to comma separated value (CSV) lists using ParaView [15]. We also use VMTK find the portions of the surface that surround each branch in a given tree; these are known as group surfaces and are also saved in CSV format using ParaView. We were unable to compute the group surface for one of the surfaces using VMTK, so this surface is neglected for the rest of the analysis ^1^. There are now 47 surfaces. The networks have between 7 and 28 terminal vessels. The group surfaces and centrelines are shown for these extrema in Fig. 4.

**Figure 4:**
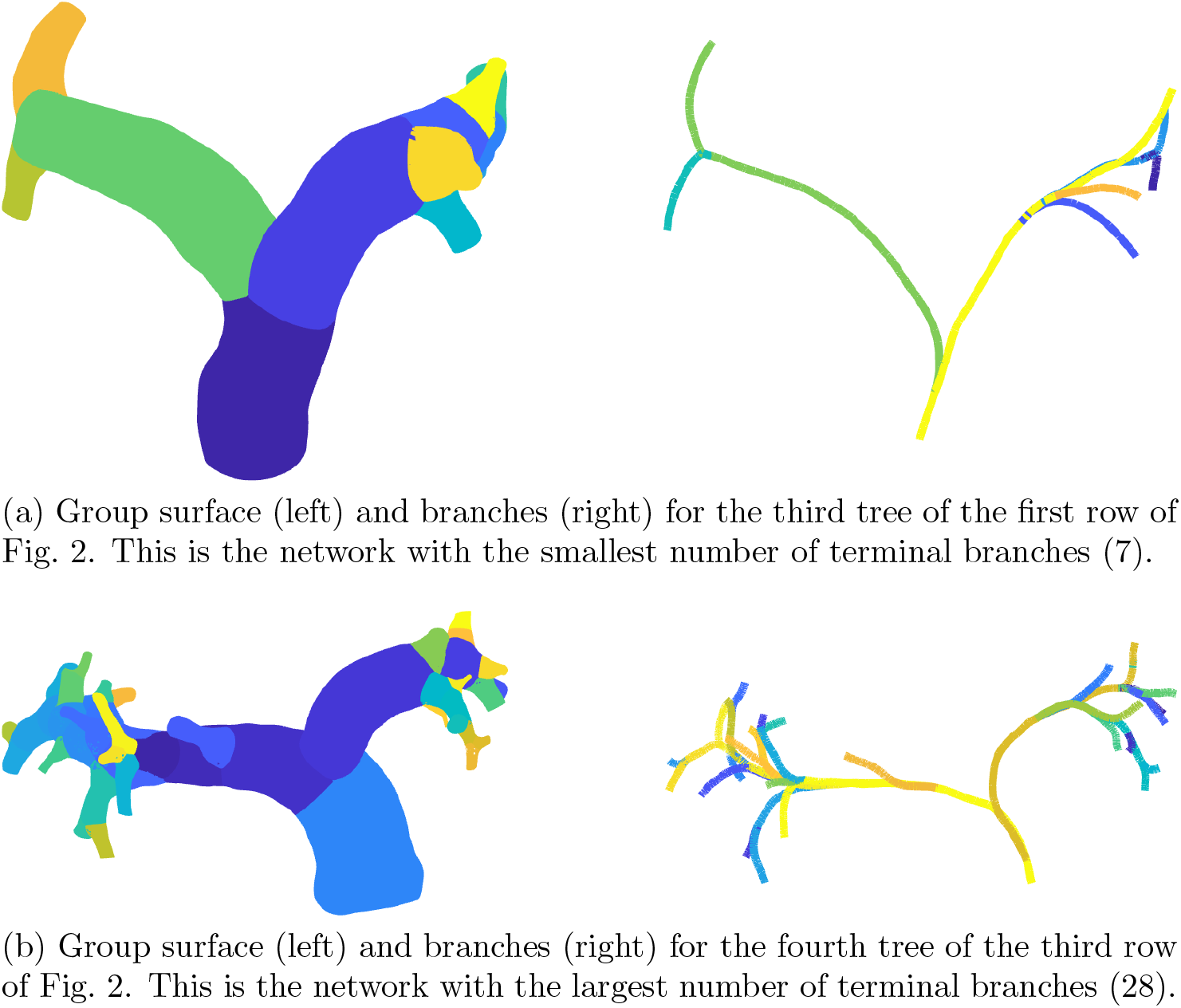
Group surfaces and branches for two of the pulmonary meshes produced using VMTK and visualised in MATLAB.

The diverging vasculatures can be represented using a graph *G* = (*V, E*) with vertices *V* and edges *E*. Since the vasculature is diverging from the inlet, and bulk flow is unidirectional, the *G* is a directed, rooted tree. The vertices *V* are the points along the centrelines with attributes of location in ℝ^3^ and a real valued radius which comes from the method of maximally inscribed spheres employed by VMTK to find the centrelines. Briefly, *G* is obtained by considering all vertices *V* that lie on the centrelines described by the centrelines and indexing these. This allows us to form edges *E* from the centrelines. Any multiply counted entries are removed from *V* and *E*. The collection of vertices *V* is now unique. Self-loops are also removed from *G*.

An edge runs between two vertices which define the centerlines. In the context of arterial graphs, it often makes more sense to think about *segments* than edges. A *segment* is a collection of edges that runs between an exterior vertex and a junction or two junction vertices. An exterior vertex is the inlet vertex (unique) or a terminal vertex – it is joined to exactly one other vertex. An exterior segment is a segment that contains an exterior vertex. A junction vertex is joined to three or more other vertices. Creating segments from edges preserves the branching structure of the graph, and all spatial properties while reducing the number of vertices and edges that need to be considered, resulting in a simpler structure to analyse. Let the collection of segments be denoted *E*^′^ and the collection of exterior and junction vertices be *V* ^′^ to form a graph *G*^′^ = (*E*^′^, *V* ^′^).

Segments are short portions of centrelines that are constructed from vertices that each have a position in space and a radius. The mean direction of a segment is that of the straight line between the initial and final vertices in a segment. Pulmonary vessels are not particularly torturous, so we are usually able to represent the surface well with straight segments. We treat the mean direction of a segment as the long axis of a cylinder that is drawn with least radius associated with the segment. Around this collection of cylinders, we can draw the original surface. This section is essentially concerned with making the collection of cylinders as similar as possible to the surface. Such a comparison between cylinders and surface can be seen in Fig. 6a.

#### 2.2.2. Data Handling and Error Correction

While VMTK produces centrelines reliably, these often require some additional correction to reflect the geometry as faithfully as possible. Regardless, VMTK is a useful tool for analysing structures of tubes and we are unaware of a more reliable tool that is as widely used. Here, we discuss the errors that arose while processing the data. These pertain to the structure of the graph, the length of segments, and the location of the junction nodes. *Twin Vessels*. The arterial network is assumed to be a tree in the graph theory sense – two vertices are connected by exactly one path. If two segments begin and end with the same vertex then they are called “twins”. Twin vessels amount to loops in the graph. We have assumed that the graph is a tree based on the gross anatomy, so such loops are not permissible. Twin segments are typically short – containing only around 4 vertices – so are difficult to see by inspecting the spatial trace of the segments. However, they are easy to see by examining the branching structure itself. An example of a network with twin vessels can be found in Fig. 5. Of 43 networks that we analyse, 16 have at least one pair of twin vessels that need to be merged. In general, we reason that we should keep the “larger” twin. In the first instance, keep the segment that contains more edges. If both twins contain the same number of segments, keep the one with larger mean radius.

**Figure 5:**
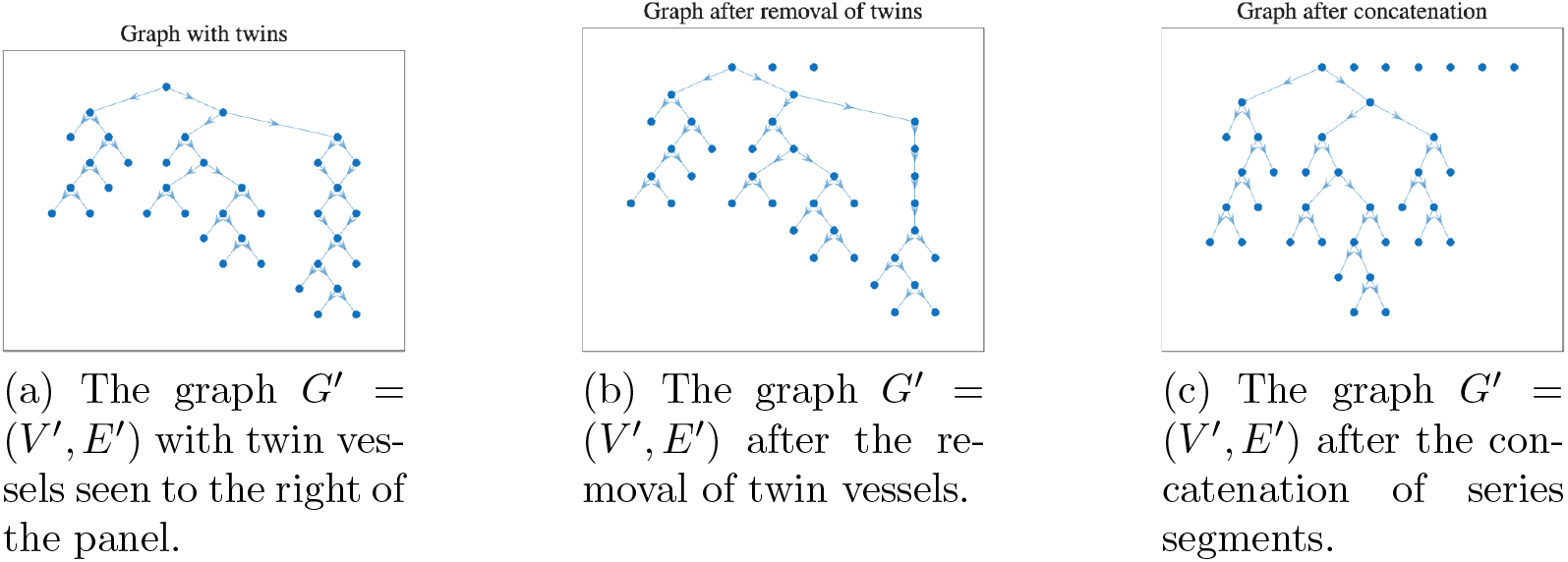
Plots of *G*^*′*^ = (*V* ^*′*^, *E*^*′*^) for a single analysed network before the removal of twin segments (a), after the removal of twin segments but before concatenation to make a single trunk (b) and after the concatenation (c). In (b) and (c) disconnected vertices can be seen. These represent the two twins from two pairs that have been removed (b) and the 4 segments that have been absorbed into a single vessel during the concatenation. The arrows along the edges *E*^*′*^ show the direction of flow from the prescribed root.

As we can see from Fig. 5, the removal of twin vessels results in two or more vessels joined in series with each other. As discussed by Mackenzie et al. [16], we concatenate such vessels to simplify the network structure without loss of spatial information. Concatenation is carried out on the basis of the number of daughters that a given segment has: if a given segment gives rise to exactly one other segment, these are concatenated.

##### Centreline Extension

As centrelines are computed in VMTK using the method of maximally inscribed spheres, all exterior segments fall short of the position that they should achieve based on the surface. This can be appreciated in Fig. 6. We are able to correct this by extrapolating the exterior segments in the direction of the exterior edge oriented towards the exterior vertex of the segment itself. The extrapolated centrelines are checked against the reference surface to ensure that all extensions that are retained lie within the surface. This process is illustrated in Fig. 6b which shows the interior segments in black, exterior segments in cyan, the accepted extensions in yellow (with rejected extensions in magenta), and the surface within which all centrelines must lie in red. We determine if a vertex belonging to a proposed segment extension lie within the surface by using a system-specific function `inpolyhedron` [17].

**Figure 6:**
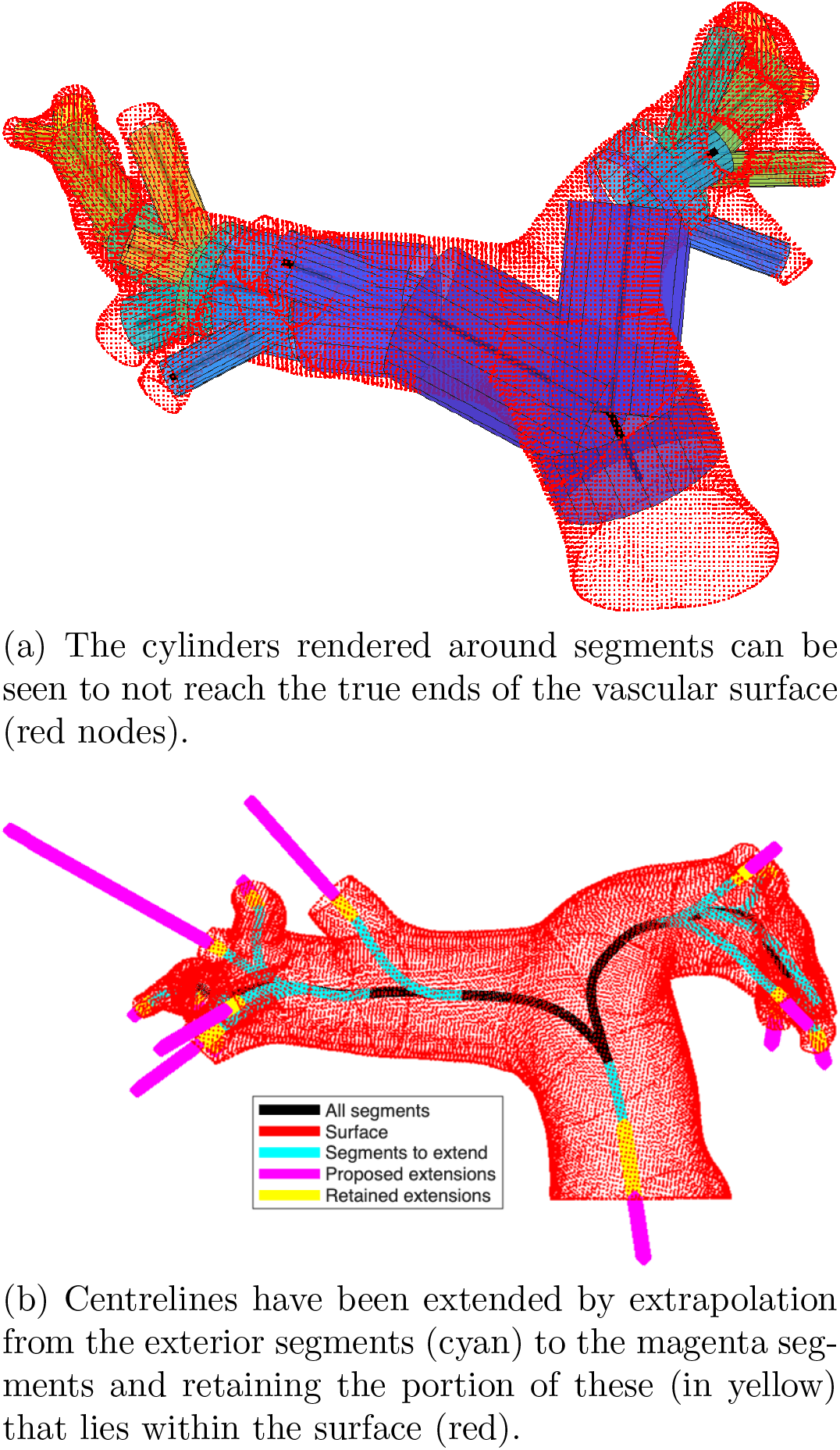
The surface and centrelines of a single pulmonary tree (the first example from Fig. 2) oriented such that the plane in which the lumen of the MPA lies is perpendicular to the page. This view highlights that the exterior segments are too short.

The extensions to the centrelines require extrapolations of the radius data so ensure that vessel radius is defined along the whole length of a vessel. To do this, for each extended segment, construct a function *r*(*d*) where *d* is the arc length from the vessel inlet and *r* is the radius. Find the change points of *r*(*d*) (using the in-built `changepoint` function in MATLAB R 2025b). For the case of the inlet segment, the line of best fit is found from the proximal end of the unextended segment until the position of the first change point. This linear function is used to extrapolate the radius backwards to the proximal end of the extended segment. For terminal segments, this process is essentially reversed. A linear function is fitted from the final change point of *r*(*d*) to the end of *r*(*d*) which is then extrapolated distally. This process is illustrated for a single representative segment in Fig. 7. This method may result in the prediction of negative radius values. If this is the case then a constant value is used as the extrapolated radius value. The constant chosen is the radius found using VMTK at the distal end of the segment.

**Figure 7:**
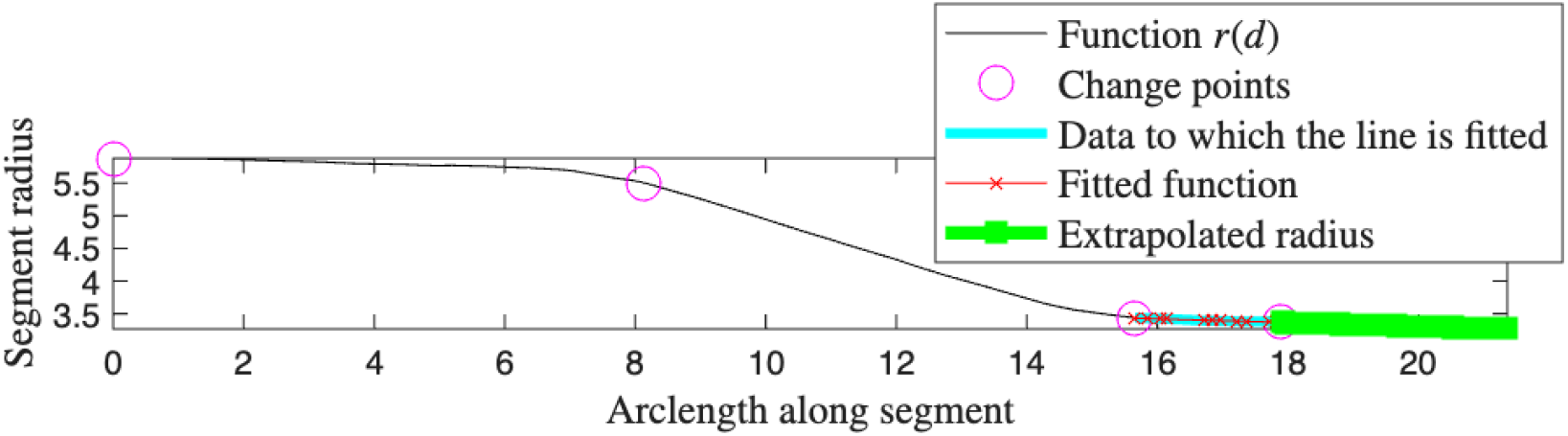
Radius extrapolation process. Since segments are extended, we require additional radius information. The radius data for the original portion of the segment is shown as the black line. We compute the change points of these data which are highlighted as magenta circles. For a terminal segment, the final portion of the radius data is used to extrapolate the radius data (cyan line) by finding the line of best fit (red crosses) and extrapolated to the segment extension (solid green).

##### Position of the junction points

By inspecting Fig. 6a we can appreciate that the bifurcation from the MPA into the left and right pulmonary arteries occurs too proximally. As discussed by Bartolo et al. [18] network morphometry (the length and radii of segments in the network) are key determinants of simulation outcome. Hence, we should endeavour to create a 1D computational domain that represents the 3D surface as faithfully as possible. If junctions between segments occur too proximally, then no segment involved in a junction can be of the correct length. Hence, we seek to shift junctions distally to more feasible locations. Bartolo et al. [18] also encounter this, and propose a method for shifting bifurcations distally based on the least Euclidean distance between nodes in a pair of daughter vessels. We propose a new method for determining where a junction should lie and extend it to 1-to-many vessel branching. This condition is based on the calibre of the vessels and the relative position of the centrelines. Essentially, if the centrelines of all the daughter segments of a single parent lie within all of its siblings for part of their length, the daughters are merged in this region and then absorbed into the parent, essentially elongating the parent and shifting the junction vertex.

For example, consider the two daughters seen in Fig. 8a. The segments are shown in solid lines and at each vertex is represented by the coloured patches. We can see that for the first third of the daughter vessels that each lies within the other. We regard this as a single vessel so the daughters should be merged in this region. Vessels are merged to the unweighted centroid in the region to merge – this is seen in cyan in Fig. 8b. Figure 8c shows the result of correcting the positions of all junction vertices in this manner. The radius at each vertex in a merged segment is computed as the maximal value of the radii of the vertices that were merged to form the current vertex. The collection of segments is traversed from the terminal segments towards the inlet segment to avoid the need to additionally pair-wise compare the daughters in a trifurcation (or a higher order junction).

**Figure 8:**
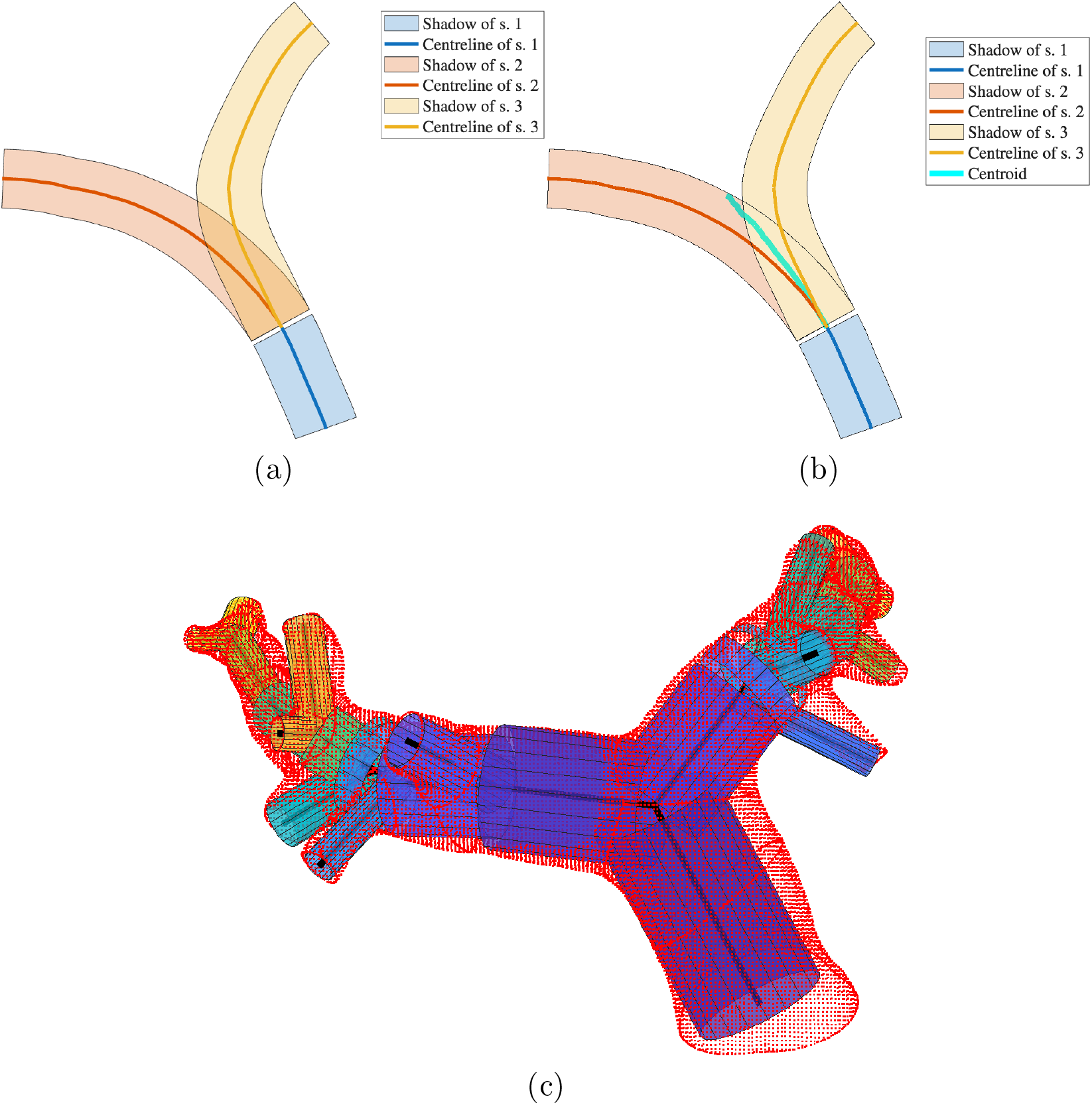
Shifting the bifurcation point of the MPA distally. (a): The centrelines of the MPA, LPA, and RPA are shown within their respective segments and the overlap of the LPA and RPA is obvious. (b) A new line shown in cyan is constructed half way between the LPA and RPA in the region in which they overlap. The new bifurcation point is chosen along this line. (c): The bifurcation points have been shifted distally and the collection of cylinders matches the net more closely than in Fig. 6a.

##### Spuriously Short Segments

A segment is considered spuriously short if the arc length of the segment is less than the minimal radius of that segment. If a short segment is removed, the tree structure is maintained. Specifically, if the *i*-th segment is removed, it’s parent becomes the parent of the daughters of the *i*-th segment. An example of a short segment can be seen in Fig. 9a and the network after the removal of the short segment can be seen in Fig. 9b.

**Figure 9:**
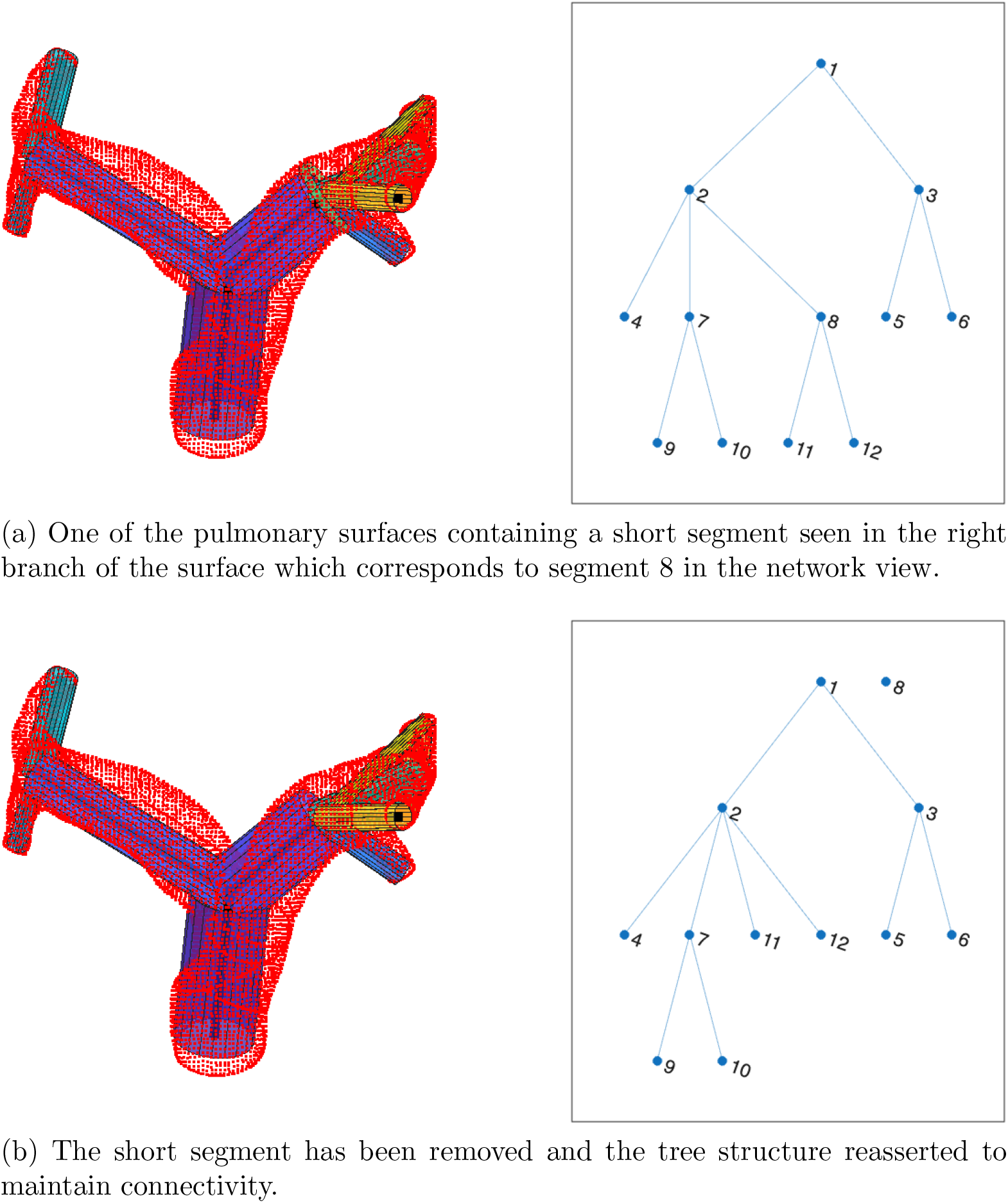
A short segment is found after the analysis discussed above and is removed before reasserting the tree structure. Segment 8 is short so is removed. The parent of segment 8 becomes the parent of the daughters of segment 8. This network can also be seen in Fig. 4a.

Segments containing two or fewer vertices are always considered short, regardless of radius, and should be dealt with before junction smoothing if they arise due to shifting bifurcation points.

##### Junction Smoothing

Shifting the junction vertices has the potential to introduces kinks into the segments which are not seen in Fig. 8c. Let *A* be the set of the *n*-many positions of the vertices of the segment each of which is in ℝ^3^. Translate all vertices in *A* so that the first vertex of the segment, *A*(1), lies at the origin. Let *B* = *A\A*(1) be the proper subset of *A* that contains all vertices of *A* that do not lie at the origin. Define *C* such that |*C*| = *n* − 1, *C*(1) lies at the origin and *C*(*n* − 1) = *B*(*n* − 1). Between the ends, let *x*_*i*_ = 1 − *i*Δ*N, i* = 0, 1, 2, …, *n* − 2, *n* − 1 where Δ*N* = 1*/*(*n* − 1) so *C*(*i*) = *B*(*i*) − *x*_*i*_*B*(1). This is a translated linear interpolation from the origin to the end of the current segment. The true segment is obtained by translating this new segment back to the original initial vertex of *A*. This procedure reduced the length of all daughter segments by 1. A similar scheme is presented by Bartolo et al. [18].

Repositioning the junction vertices changes the lengths of the segments. Some vessels may become so short that they are absorbed into neighbouring vessels to avoid violating the modelling assumption that the length of a segment is much greater than its radius. Repositioning the junction points may also result in an entire segment being absorbed into its parent. In such cases, it is important to reassert the tree structure so that no disconnected components are created.

#### 2.2.3. Creation of computational domains

##### Removal of downstream vessels

Here, we are interested in simulating blood flow in the pulmonary circulation after lung resection. We have now constructed 1D representations of 45 different pulmonary vascular networks. Three networks could not be analysed due to errors in the centrelines constructed with VMTK. Since each vessel in all of the networks are discrete mathematical objects, it is straight forward to systematically remove any given vessel and all downstream vessels. For a given network, initially remove in turn any vessel (and downstream vessels) that are not the MPA, left pulmonary artery (LPA), or right pulmonary artery (RPA) trunks. A new geometry is created for each alteration to the original network. These alterations are illustrated in Fig. 10 on the network shown in Fig. 9. Here, the removed vessel is highlighted in cyan and consequently removed downstream vessels in magenta. The networks are shown in order of decreasing volume. The MPA is never removed. The unaltered network appears in the top left panel.

**Figure 10:**
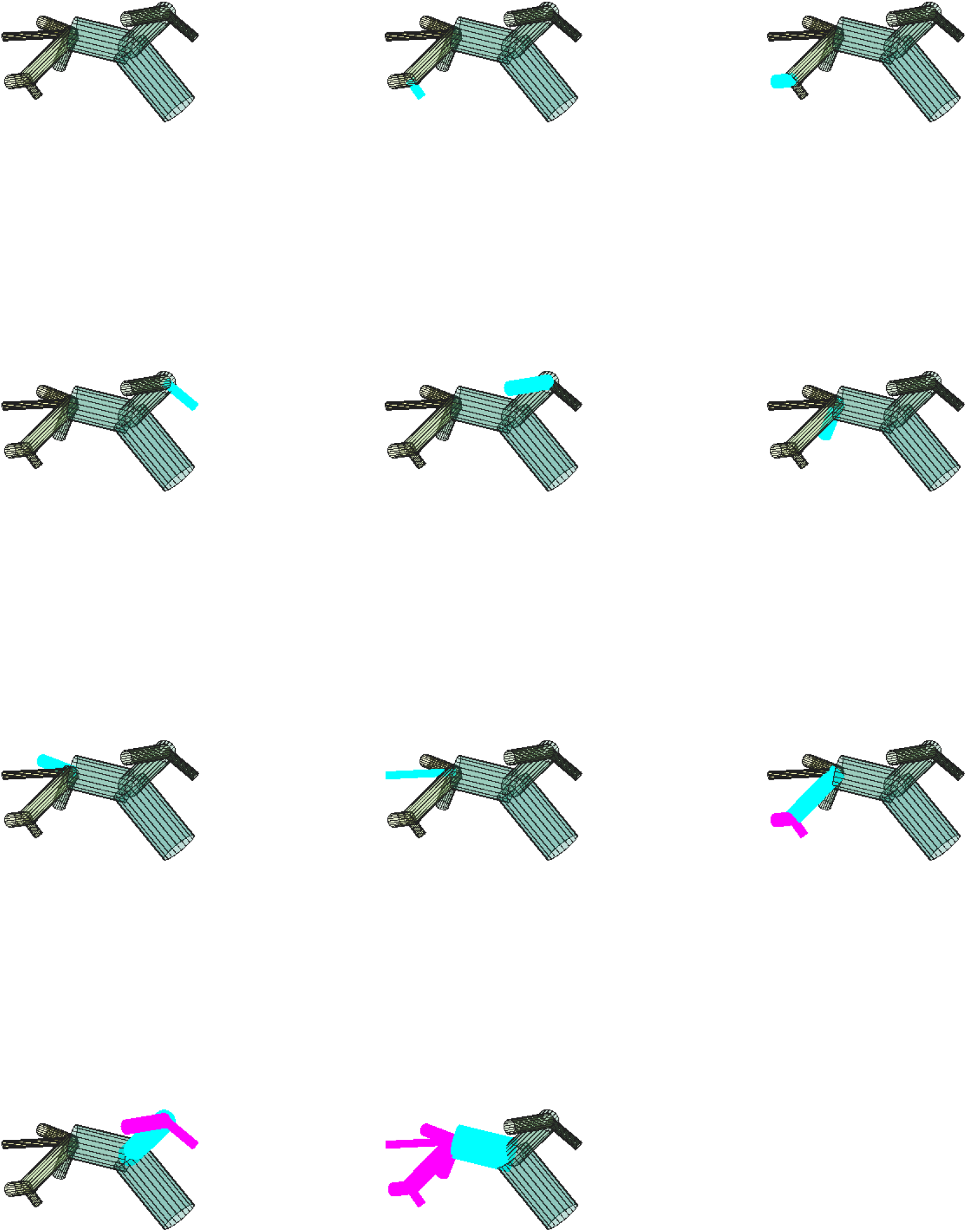
Exemplar of vessel removal from the network seen in Fig. 9. The removed vessel is highlighted in cyan. The downstream vessels in magenta are also removed. The view differs from that of Fig. 9 so all vessels can be appreciated. The networks are arranged in order of volume post vessel resection.

Additionally, combinations of terminal vessels are removed to create new altered networks. Two (or more) terminal segments that are siblings are removed together with and without any aunts. We choose to make these alterations as they represent larger resections than removing single terminal vessels, and are thus more likely to be clinically realistic.

### 2.3. Flow simulations

Of the original networks, 5 networks have bifurcations only, 27 have at least one trifurcation, 12 with at least one junction with one-to-four branching, and a single tree with a five-to-one junction. The network with one-to-five branching is excluded from further analysis. There are 44 unaltered networks in which we simulate flow. Of these 44 networks, we make 1602 alterations so there are 1646 networks in total.

We create tabular representations of the original and altered networks which capture their morphometric properties: the length and radius of each branch in each network including the connectivity of each vessel. Segments are modelled as straight walled cylinders in which the proximal and distal radii are set as the mean radius of the segment. In all networks, the initial vessel is the MPA, the first branch from this is the LPA and the second branch is the RPA.

We simulate flow in all 1646 networks using the boundary conditions as described above. The simulations run until convergence or a failure to converge. Convergence is checked by comparing flow in two successive periods. Let 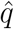 be flow at the distal end of a single vessel in the current period and *q* be the same quantity but in the previous period. Then define SSE as 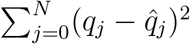. Compute this at the distal end of all vessels for all periods. The simulation is deemed to have converged when the SSE falls below 1 in all vessels. Simulations are given 20 iterations in which to converge. In total, 6 simulations failed to converge. No simulations in unaltered networks failed. There are results from 1640 simulations to analyse. From these, we save fluid pressure, volumetric flux, and lumen area at all spatial points (every 0.2 cm) in the MPA, LPA, and RPA for a whole cardiac cycle after the computational scheme has converged. Simulations take an average of around 5 s of computational time to converge^2^ for a total computational time of around 2 hours.

### 2.4. Analysis of Simulation Results

So far, we have produced representations pulmonary arterial trees and alterations thereof in which we have simulated blood flow. We now analyse the simulated flow data to determine the impact of lung resection on pulmonary arterial pressure waves. We are interested in a simulated reproduction of the findings reported by Glass [8] and Glass et al. [6]. In particular, we seek to investigate whether we can simulate the change in wave intensity in the altered trees as compared to the unaltered tree. This investigation is undertaken as a computational model that can predict increased right ventricular load via wave intensity has the potential to be clinically useful as a treatment-planning tool.

Glass et al. [6] analyse pre and post-operative MRI images in 27 lung resection patients. From these images, they are able to derive volumetric flux of blood *q* and lumen area *A* in the right and left pulmonary arteries (where present). They smooth the data using a 7-point, second-order Savitzky-Golay filter [19]. The data are then interpolated to 1 ms using a cubic spline without further smoothing. From these data they compute wave intensity for each patient in both the left and right PAs pre- and post operatively following Quail et al. [20].

### 2.5. Computing Wave Intensity

In this study, we seek to replicate the findings pertaining of Glass [8] and Glass et al. [6] with a computational model. The main findings of Glass and colleagues pertain to changes in wave intensity post-lung resection. To compare the output of our computational model against their measured data, we must compute wave intensity.

Wave intensity is a measure of the power transmitted by the pulse wave as the waves hit the internal walls of the vasculature. From the simulation results we have pressure *p, q, A* and we can compute fluid velocity *u* = *q/A* and pulse wave velocity

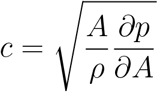

in which *ρ* = 1.055 kgL^−1^ is fluid density of blood. There are many alternative definitions of pulse wave velocity. We choose this one as it is easy to compute and, other than *ρ*, needs no information other than the simulated data.

Essentially, we wish to separate the pulse wave into forwards propagating and reflected components. To do this, find the differences

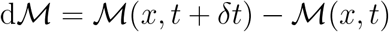

for ℳ = *q, A, p, u* where *δt* = 1 ms is the time-step size. These differences dℳ are separated into the forward and backwards propagating components dℳ_±_ such that dℳ = dℳ_+_ + dℳ_−_. As shown by Parker [21], these differences together with the water hammer equation for forwards and backwards waves

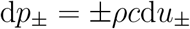

can be solved for changes in the forwards and backwards waves

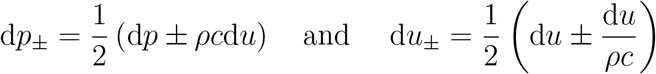

giving

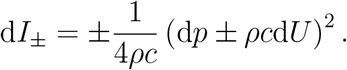

This is the definition of wave intensity adopted by, for example, Qureshi & Hill [22] and Lechuga [23].

There is an alternative definition of wave intensity d*I*_±_ = d*Q*_±_d*A*_±_ in which

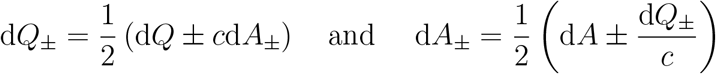

from *c* = ±d*q*_±_*/*d*A*_±_ yielding

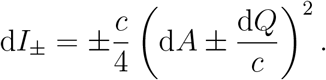

Here, d*I*_±_ are the forwards and backwards propagating components of the pulse wave. As can be seen in Fig. 11, the two definitions of wave intensity yield very similar time series up to a scale factor with units kg m^−3^ s^−2^. The units and magnitude of the peaks are the major differences between the curves. From this figure, we note that the wave intensities as comparable to those found in the literature [6, 8, 20, 22, 23, 24].

**Figure 11:**
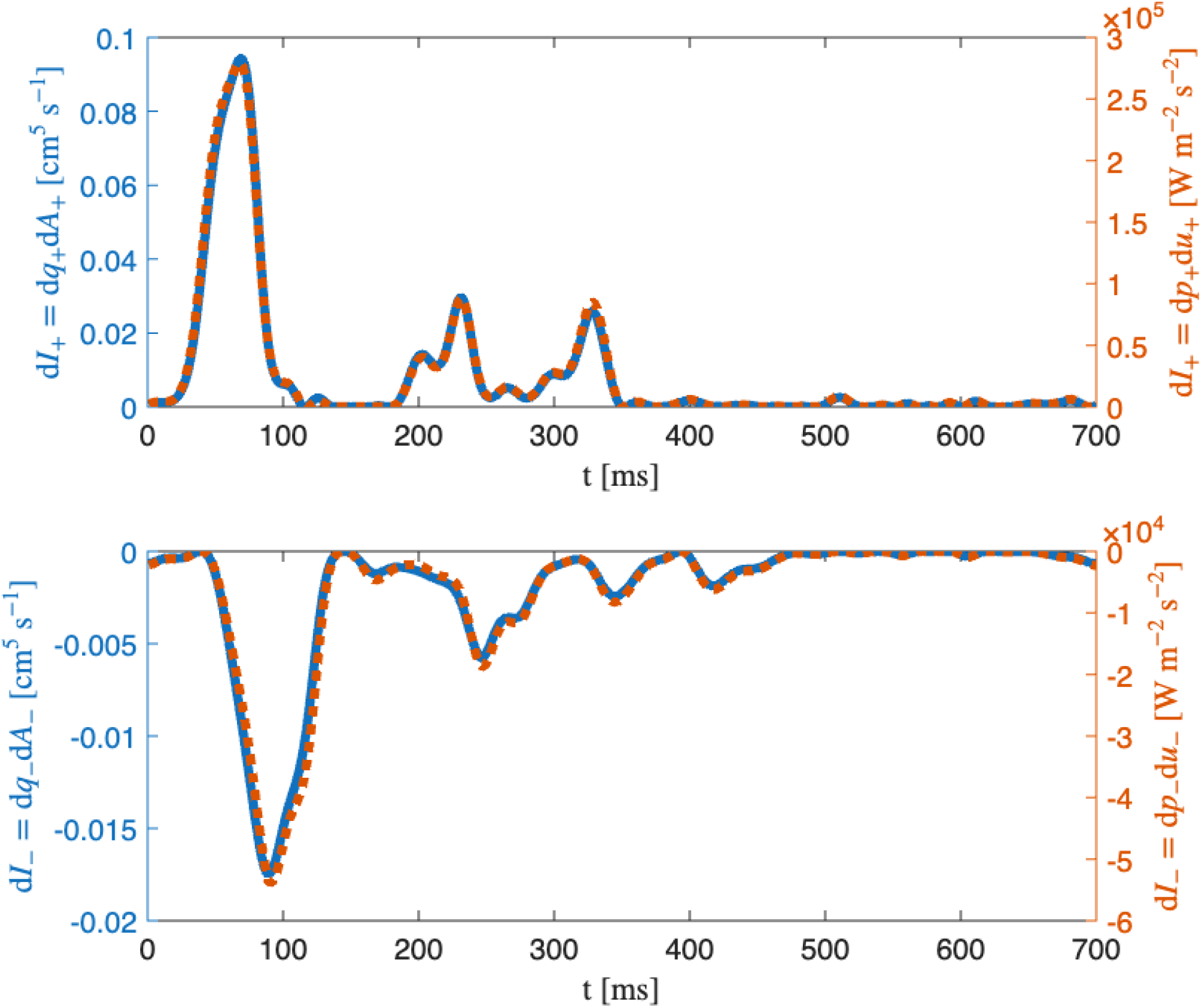
A comparison between the wave intensity time series when computed as d*I*_*±*_ = d*q*_*±*_d*A*_*±*_ (solid blue, left hand scale) and d*I*_*±*_ = d*p*_*±*_d*u*_*±*_ (orange dotted, right hand scale).

As highlighted by Parker [21], the sampling period (the time step size) will impact the computed wave intensity. To avoid issues when comparing the simulated flows against the measured flows of Glass, we have chosen the same sampling period. Alternatively, we could have used the time-step invariant definition of wave intensity d*I*^′^ = d*I/*(*δt*^2^) as seen in Qureshi & Hill [22].

### 2.6. Data Interpolation

We adopt the same wave intensity definition as Glass et al. [6] to simplify further comparisons. As we have adopted the same wave intensity definition, which does not take time-step size into account, we must also use a time step size of 1 ms. We achieve this by linearly interpolating the simulated data from 512 time steps per 700 ms period to 701 time-steps before computing wave intensities.

Further, we also apply a Savitzky-Golay filter to the data. This filtration method smooths data while retaining the signal tendency. The filter is applied after interpolation to time-steps of size 1 ms. We apply it to three successive periods of *p, q*, and *A* for all simulations. Three periods are necessary as the application of the filter to a single period may incur a mismatch between the two ends of the period. When applying the filter, we must choose the window length over which it is applied. Choose window length to be the largest value that gives an SSE of less than 1 between the filtered and unfiltered signals in the middle of the 3 successive periods. Window range is between 13 and 31 with a median length of 19 and 98.72% of window lengths being between 17 and 21.

#### 2.6.1. Wave components

The time series d*I*_±_ are forwards and backwards propagating waves. From each of these we wish to isolate the compressive and decompressive parts in order to analyse the different wavefronts separately. Decompressive waves are sometimes called expansion waves in the literature [7]. There are four wave components:

- Forward compression wave (FCW): d*I*_+_ *>* 0 and d*p*_+_ *>* 0,
- Forward decompression wave (FDCW): d*I*_+_ *>* 0 and d*p*_+_ *<* 0,
- Backward compression wave (BCW): d*I*_−_ *>* 0 and d*p*_−_ *>* 0,
- Backward decompression wave (BDCW): d*I*_−_ *>* 0 and d*p*_−_ *<* 0.

These four distinct wave components can be appreciated in Fig. 12.

**Figure 12:**
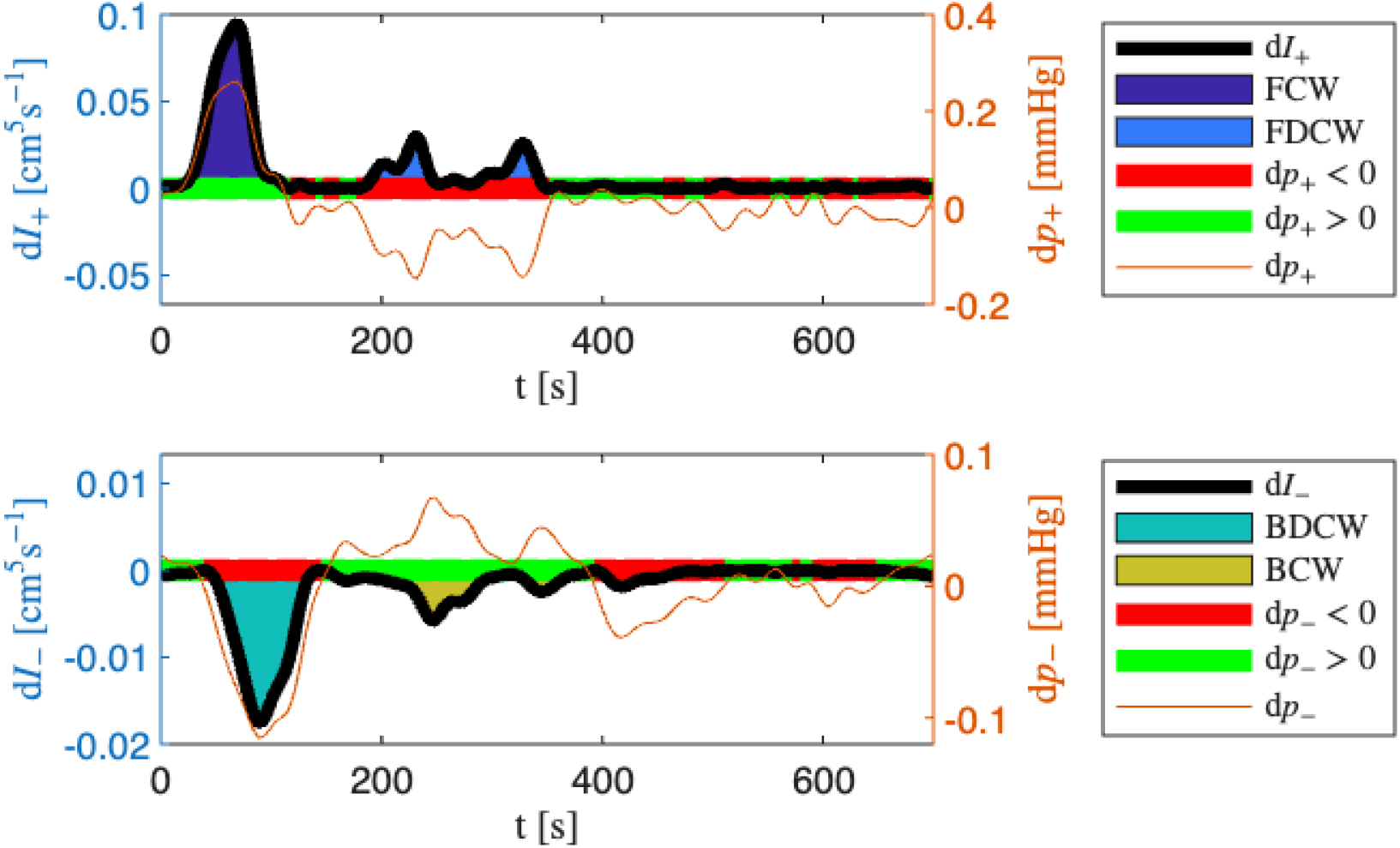
Forwards and backwards waves are separated into compressive and decompressive regions based on the sign of d*p*_*±*_. The *t*-axis is highlighted green when d*p*_*±*_ *>* 0 (compression) and red when d*p*_*±*_ *<* 0 (decompression).

Let

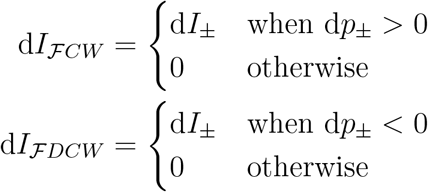

in which ℱ = *F* (forward waves) when considering d*I*_+_ and ℱ = *B* (backward waves) when considering d*I*_−_. There are three parameters of interest that we investigate for each of these four wave components. The first is peak wave intensity value defined as 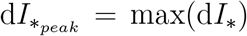 for ∗ = *FCW, FDCW* and 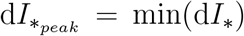 for ∗ = *BCW, BDCW* . The second is the first time at which this occurs *t*_*peak*_. The third is the absolute value of the integral 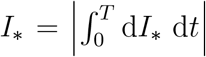 over a single period of length *T* = 700 ms for ∗ ∈ {FCW, FDCW, BCW, BDCW}. We compute these for all completed simulations at the distal ends of the MPA, LPA, and RPA. These are the data that will be analysed in the upcoming sections. We have chosen these in order to compare our results against those presented by Glass [8].

## 3. Results

Glass [8] examines the three parameters of interest for the three wave components FCW, FDCW, and BCW but neglects the BDCW. As such, we also neglect BDCW as there is no data against which to compare our findings. We wish to compare the three parameters of interest for the three wave components between the operative and non-operative pulmonary arteries pre- and post-operatively that are the output of our simulations and processing steps. The unaltered pulmonary tree is considered to be the pre-operative state. In cases of simulated lobectomy, the operative pulmonary artery is either the LPA or RPA that lies upstream of the vessels removed in a given tree.

### 3.1. Pre-operative wave intensity in the MPA, LPA, and RPA

As can be seen in Figs. 2 & 3 the pulmonary circulatory system is not symmetrical in the right and left sides. As such, we should not expect that the wave intensities in the right and left sides should not behave identically, especially in cases of unilateral changes to the pulmonary artery geometries. We tabulate the parameters of interest in each of the four wave components in each of the MPA, LPA, and RPA in Table 1.

**Table 1:** Comparison between *t*_*peak*_, d*I*_*peak*_ and *I*_*W*_ for each of the 3 major pulmonary arteries and each of the 4 wave components. These values are given as mean (standard deviation) to 4 decimal places. *n* = 44.

| Wave | Parameter | MPA | LPA | RPA | Unit |
| --- | --- | --- | --- | --- | --- |
| FCW | $t_{peak}$ | 67.1818 (4.8189) | 66.9091 (14.3362) | 59.6591 (15.4498) | ms |
| | $dI_{peak}$ | 0.1053 (0.0178) | 0.0402 (0.0108) | 0.0449 (0.0126) | $\text{cm}^5 \text{s}^{-1}$ |
| | $I_W$ | 4.4026 (0.6965) | 1.6546 (0.4646) | 1.8106 (0.5045) | $10^{-3}\text{cm}^5$ |
| FDCW | $t_{peak}$ | 273.3636 (49.7300) | 304.5000 (48.6795) | 293.1591 (48.5041) | ms |
| | $dI_{peak}$ | 0.0292 (0.0036) | 0.0106 (0.0032) | 0.0116 (0.0032) | $\text{cm}^5 \text{s}^{-1}$ |
| | $I_W$ | 2.0658 (0.2145) | 0.7559 (0.2425) | 0.8287 (0.2533) | $10^{-3}\text{cm}^5$ |
| BCW | $t_{peak}$ | 229.5227 (26.3381) | 182.5909 (44.6085) | 186.3182 (91.6413) | ms |
| | $dI_{peak}$ | -0.0047 (0.0015) | -0.0015 (0.0007) | -0.0017 (0.0015) | $\text{cm}^5 \text{s}^{-1}$ |
| | $I_W$ | 0.3913 (0.0948) | 0.1003 (0.0486) | 0.1219 (0.0743) | $10^{-3}\text{cm}^5$ |
| BDCW | $t_{peak}$ | 103.9773 (71.0331) | 174.0000 (136.3064) | 152.7955 (135.3022) | ms |
| | $dI_{peak}$ | -0.0119 (0.0063) | -0.0025 (0.0023) | -0.0039 (0.0034) | $\text{cm}^5 \text{s}^{-1}$ |
| | $I_W$ | 0.6487 (0.2870) | 0.1239 (0.0909) | 0.1782 (0.1376) | $10^{-3}\text{cm}^5$ |

For each unaltered network (*n* = 44), each pair of pulmonary arteries (MPA, LPA, and RPA), for each wave component, and for each parameter of interest, we are able to assess whether the distributions of values exhibit a statistically significant difference in mean value. We assess this by carrying out a paired Student’s *t*-test. We choose this test as the data are paired samples. Because we are only looking at the unaltered trees, *n* = 44. The significance of the difference between the distributions for the parameters of interest between the major vessels is seen in Table 1 is summarised in Table 2. The null hypothesis is that the distributions have the same mean value. The null hypothesis is accepted for all parameters compared between the LPA and RPA except for *t*_*peak*_ in the FCW and d*I*_*peak*_ in the BDCW. This indicates that flow to each lung behaves similarly in the pre-operative cases. For comparisons between the MPA and either daughter, the null hypothesis is rejected for all parameters of interest and for all wave components except for *t*_*peak*_ between the MPA and LPA in the FCW.

**Table 2:**
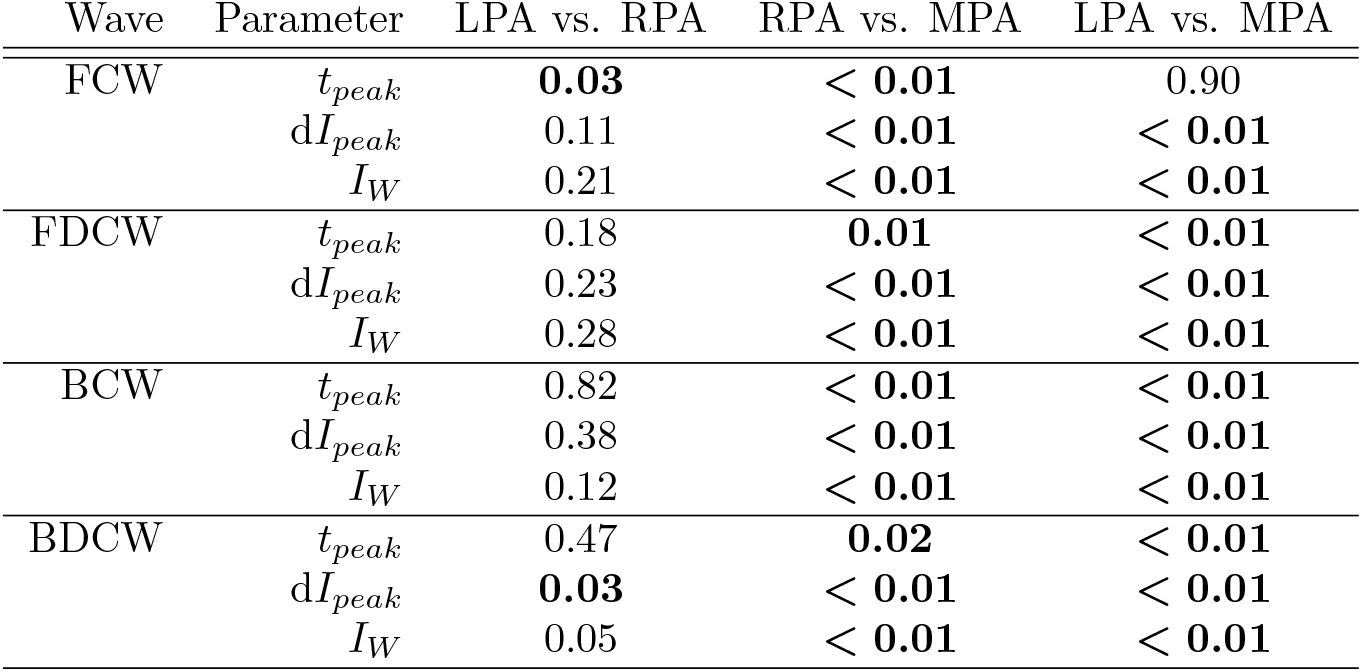
*p*-values obtained by pair-wise comparisons of the distributions of the parameters of interest between the major blood vessels using a paired-sample *t*-test. The significance level is chosen as *p <* 0.05. Significant results are emboldened. If *p <* 0.01, the value is not given other than as the inequality.

| Wave | Parameter | LPA vs. RPA | RPA vs. MPA | LPA vs. MPA |
| --- | --- | --- | --- | --- |
| FCW | $t_{peak}$ | **0.03** | $< \mathbf{0.01}$ | 0.90 |
| | $dI_{peak}$ | 0.11 | $< \mathbf{0.01}$ | $< \mathbf{0.01}$ |
| | $I_W$ | 0.21 | $< \mathbf{0.01}$ | $< \mathbf{0.01}$ |
| FDCW | $t_{peak}$ | 0.18 | **0.01** | $< \mathbf{0.01}$ |
| | $dI_{peak}$ | 0.23 | $< \mathbf{0.01}$ | $< \mathbf{0.01}$ |
| | $I_W$ | 0.28 | $< \mathbf{0.01}$ | $< \mathbf{0.01}$ |
| BCW | $t_{peak}$ | 0.82 | $< \mathbf{0.01}$ | $< \mathbf{0.01}$ |
| | $dI_{peak}$ | 0.38 | $< \mathbf{0.01}$ | $< \mathbf{0.01}$ |
| | $I_W$ | 0.12 | $< \mathbf{0.01}$ | $< \mathbf{0.01}$ |
| BDCW | $t_{peak}$ | 0.47 | **0.02** | $< \mathbf{0.01}$ |
| | $dI_{peak}$ | **0.03** | $< \mathbf{0.01}$ | $< \mathbf{0.01}$ |
| | $I_W$ | 0.05 | $< \mathbf{0.01}$ | $< \mathbf{0.01}$ |

### 3.2. Post-operative comparisons

We are interested in whether the computational framework described above can capture the qualitative behaviour of the experimental results described by Glass [8]. Glass produces one table and one box-plot chart per wave component. Glass computes the three parameters of interest for the four wave components pre-operatively, on post-operative day 2, and at two months post-operatively. From these tables we find the post-operative change in the operative and non-operative PAs in the parameters of interest in the FCW, FDCW, BCW and summarise these changes in Table 3. There is no growth and remodelling in the computational model, so we do not compare the simulation results against the measured data from two months post-operatively. For any simulation in an altered network, we are able to determine the operative side by comparison between the current tree and the unaltered tree for a given patient. We are interested in the changes to the parameters of interest in the LPA and RPA, so we omit cases of pneumonectomy from this analysis.

**Table 3:** Table of trends for the FCW adapted from Table 7-3 as seen in Glass [8]. The column headed Δ records how a given value changes from the pre-op case. An upwards arrow (↓) is used to denote an increase; a downwards arrow (↑) is used to denote a decrease. The pattern of changes is shown for the three parameters of interest in the non-operative pulmonary arteries (N.Op.) and the operative pulmonary arteries (Op.). When the post-operative value is lower than the pre-operative value, this is a decrease from the pre-operative value. Further, we find the ratio of post-operative:pre-operative value. If the ratio exceeds 1.5, the change is deemed to be large and is denoted with a double arrow (⇈ for large increases; ↓↓ for large decreases).

| Wave | Parameter | Pulmonary artery | $\Delta$ |
| --- | --- | --- | --- |
| FCW | $t_{peak}$ | N.Op. | ↓ |
| | | Op. | ↓ |
| | $di_{peak}$ | N.Op. | $\uparrow\uparrow$ |
| | | Op. | ↑ |
| | $I_W$ | N.Op. | $\uparrow\uparrow$ |
| | | Op. | ↓ |
| FDCW | $t_{peak}$ | N.Op. | ↑ |
| | | Op. | ↑ |
| | $di_{peak}$ | N.Op. | $\uparrow\uparrow$ |
| | | Op. | ↓ |
| | $I_W$ | N.Op. | $\uparrow\uparrow$ |
| | | Op. | ↑ |
| BCW | $t_{peak}$ | N.Op. | ↓ |
| | | Op. | ↓ |
| | $di_{peak}$ | N.Op. | ↓ |
| | | Op. | $\downarrow\downarrow$ |
| | $I_W$ | N.Op. | ↑ |
| | | Op. | $\uparrow\uparrow$ |

For each admissible alteration on for each pulmonary tree we compute d*I*_±_ and d*p*_±_ in the LPA and RPA. With these, separate out the four wave components and compute the three parameters of interest for each. We are interested in the difference from the pre-operative cases, so for each simulation and parameter of interest, find the change from the pre-operative case. Further, determine if, for a given simulation, whether the removed vessels lay down stream of the LPA; if so, the LPA is the operative side. Otherwise the RPA is the operative side. Record the change in the 3 parameters of interest for each of the 4 wave components in the LPA and RPA in all cases in which the LPA was the operative side. In these cases, the RPA is the non-operative side. Similarly, record these same data in the both PAs in cases where the RPA is the operative side. These post-operative change is recorded in Table 5. Glass [8] presents data on the change of the parameters of interest for the FCW, FDCW, BCW but not the BDCW. Hence, we are able to compare the qualitative simulated change against the FCW, FDCW, and BCW. In Table 5 we highlight a cell in green if the direction of the change aligns with that reported by Glass. Of the 36 changes given, 29 are in the same direction as those reported by Glass. Of these aligned changes, 28 are statistically significant using a paired *t*-test with a significance threshold of *p <* 0.05.

The alignment of these computational and experimental results in a large collection of non-patient specific models is good evidence that the experimentally observed changes to wave intensity are due to changes in wave reflections within the PAs. Further patient specific modelling is required in order to refine this association between lung resection, wave reflections, and right ventricular injury.

In general, we expect there to be more branches of the RPA than the LPA, as the right lung is larger than the left. However, this does not appear to be reflected in the data as the number of branches that appear on each side is a function of the original segmentation. The LPA was the operative side in *n* = 823 cases and the RPA was the operative PA in *n* = 688 cases.

In Table 2, we noted that the pre-operative difference between the parameters of interest between the LPA and RPA was rarely statistically significant. This does not hold post operatively. For Table 4, we collect the three parameters of interest for each of the 4 wave components for simulations in which both the LPA and RPA are present. Using a paired *t*-test, we compare the distributions for each of these 12 parameters between the LPA & RPA, MPA & RPA, and MPA & LPA. Post-operatively, we see that the difference between the distributions of almost all parameters of interest has become statistically significant. The exception is d*I*_*peak*_ for the BCW which is approaching significance. This is to be expected as a unilateral change should induce different behaviour between the sides. This is further evidence that the model is capturing the post-operative flow redistribution.

**Table 4:** *p*-values obtained by pair-wise comparisons of the distributions of the parameters of interest between the major blood vessels using a paired-sample *t*-test. The significance level is chosen as *p <* 0.05. Significant results are emboldened. If *p <* 0.01, the value is not given other than as the inequality. Values are given to 2d.p..

| Wave | Parameter | LPA vs. RPA | RPA vs. MPA | LPA vs. MPA |
| --- | --- | --- | --- | --- |
| FCW | $t_{peak}$ | $\ll \mathbf{0.001}$ | $\ll \mathbf{0.001}$ | 0.87 |
| | $dI_{peak}$ | $\ll \mathbf{0.001}$ | $\ll \mathbf{0.001}$ | $\ll \mathbf{0.001}$ |
| | $I_W$ | $\ll \mathbf{0.001}$ | $\ll \mathbf{0.001}$ | $\ll \mathbf{0.001}$ |
| FDCW | $t_{peak}$ | $\ll \mathbf{0.001}$ | $\ll \mathbf{0.001}$ | $\ll \mathbf{0.001}$ |
| | $dI_{peak}$ | $\ll \mathbf{0.001}$ | $\ll \mathbf{0.001}$ | $\ll \mathbf{0.001}$ |
| | $I_W$ | $\ll \mathbf{0.001}$ | $\ll \mathbf{0.001}$ | $\ll \mathbf{0.001}$ |
| BCW | $t_{peak}$ | $\ll \mathbf{0.001}$ | $\ll \mathbf{0.001}$ | $\ll \mathbf{0.001}$ |
| | $dI_{peak}$ | 0.07 | $\ll \mathbf{0.001}$ | $\ll \mathbf{0.001}$ |
| | $I_W$ | $\ll \mathbf{0.001}$ | $\ll \mathbf{0.001}$ | $\ll \mathbf{0.001}$ |
| BDCW | $t_{peak}$ | $\ll \mathbf{0.001}$ | $\ll \mathbf{0.001}$ | $\ll \mathbf{0.001}$ |
| | $dI_{peak}$ | $\ll \mathbf{0.001}$ | $\ll \mathbf{0.001}$ | $\ll \mathbf{0.001}$ |
| | $I_W$ | $\ll \mathbf{0.001}$ | $\ll \mathbf{0.001}$ | $\ll \mathbf{0.001}$ |

**Table 5:**
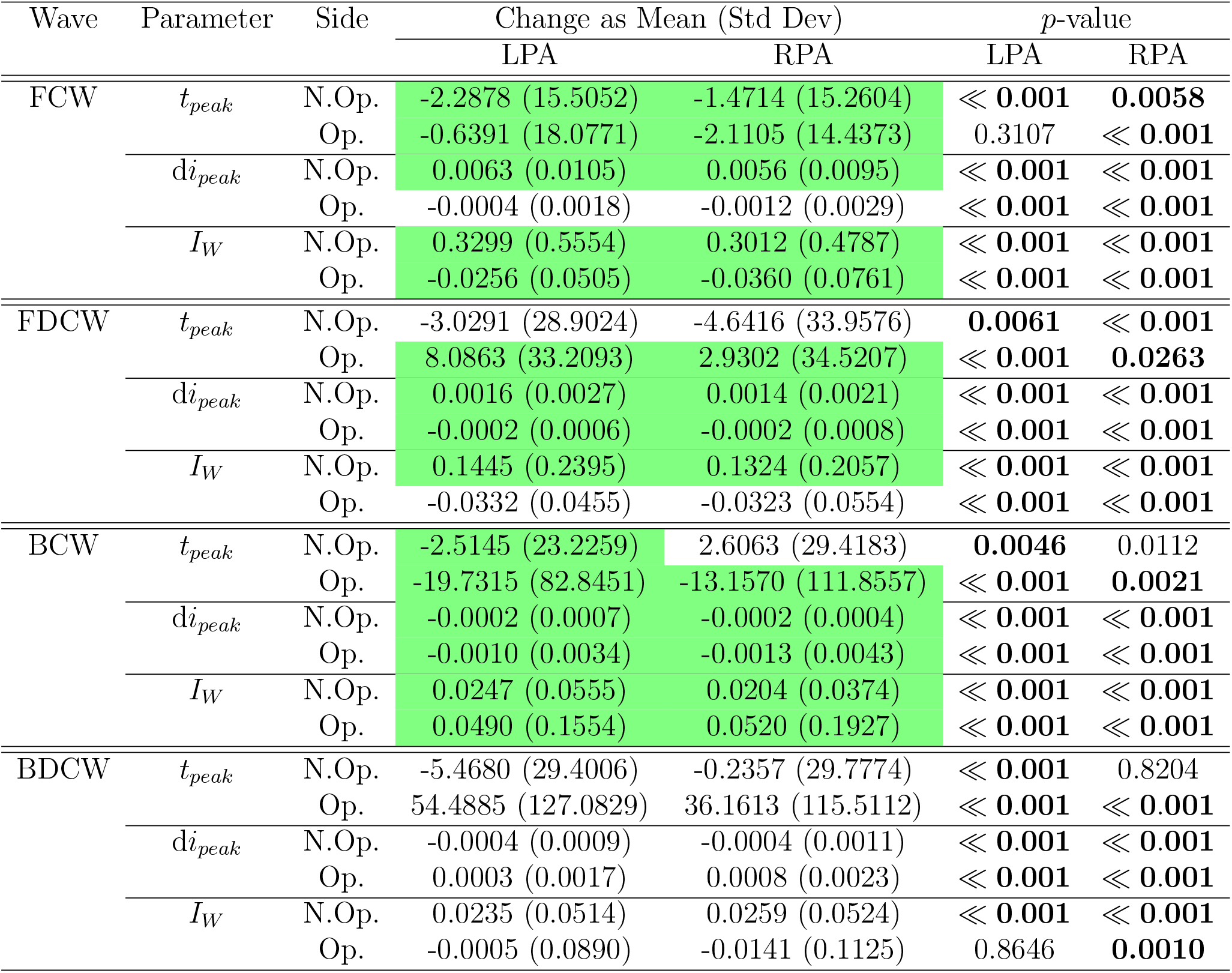
Comparisons between pre- and post operative parameters of interest in the operative and non-operative LPA and RPA before and after simulated lung resection. For each patient, for each simulation compute the change in the 12 parameters of interest as compared to the pre-operative case. A negative value means the post-operative value is lower than the pre-operative value. For the FCW, FDCW, BCW, the cells are highlighted in green if the direction of change agrees with that found by Glass [8] and summarised in Table 3. The statistical significance of each change from the pre-operative value is given in the final column. The significance threshold is chosen as *p <* 0.05. Values that meet this threshold are emboldened for emphasis. Very small values *<* 0.0001 are given as ≪ 0.001.

| Wave | Parameter | Side | Change as Mean (Std Dev) | | $p$-value | |
| --- | --- | --- | --- | --- | --- | --- |
|  |  |  | LPA | RPA | LPA | RPA |
| FCW | $t_{peak}$ | N.Op. | -2.2878 (15.5052) | -1.4714 (15.2604) | ≪ **0.001** | **0.0058** |
| | | Op. | -0.6391 (18.0771) | -2.1105 (14.4373) | 0.3107 | ≪ **0.001** |
| | $di_{peak}$ | N.Op. | 0.0063 (0.0105) | 0.0056 (0.0095) | ≪ **0.001** | ≪ **0.001** |
| | | Op. | -0.0004 (0.0018) | -0.0012 (0.0029) | ≪ **0.001** | ≪ **0.001** |
| | $I_W$ | N.Op. | 0.3299 (0.5554) | 0.3012 (0.4787) | ≪ **0.001** | ≪ **0.001** |
| | | Op. | -0.0256 (0.0505) | -0.0360 (0.0761) | ≪ **0.001** | ≪ **0.001** |
| FDCW | $t_{peak}$ | N.Op. | -3.0291 (28.9024) | -4.6416 (33.9576) | **0.0061** | ≪ **0.001** |
| | | Op. | 8.0863 (33.2093) | 2.9302 (34.5207) | ≪ **0.001** | **0.0263** |
| | $di_{peak}$ | N.Op. | 0.0016 (0.0027) | 0.0014 (0.0021) | ≪ **0.001** | ≪ **0.001** |
| | | Op. | -0.0002 (0.0006) | -0.0002 (0.0008) | ≪ **0.001** | ≪ **0.001** |
| | $I_W$ | N.Op. | 0.1445 (0.2395) | 0.1324 (0.2057) | ≪ **0.001** | ≪ **0.001** |
| | | Op. | -0.0332 (0.0455) | -0.0323 (0.0554) | ≪ **0.001** | ≪ **0.001** |
| BCW | $t_{peak}$ | N.Op. | -2.5145 (23.2259) | 2.6063 (29.4183) | **0.0046** | 0.0112 |
| | | Op. | -19.7315 (82.8451) | -13.1570 (111.8557) | ≪ **0.001** | **0.0021** |
| | $di_{peak}$ | N.Op. | -0.0002 (0.0007) | -0.0002 (0.0004) | ≪ **0.001** | ≪ **0.001** |
| | | Op. | -0.0010 (0.0034) | -0.0013 (0.0043) | ≪ **0.001** | ≪ **0.001** |
| | $I_W$ | N.Op. | 0.0247 (0.0555) | 0.0204 (0.0374) | ≪ **0.001** | ≪ **0.001** |
| | | Op. | 0.0490 (0.1554) | 0.0520 (0.1927) | ≪ **0.001** | ≪ **0.001** |
| BDCW | $t_{peak}$ | N.Op. | -5.4680 (29.4006) | -0.2357 (29.7774) | ≪ **0.001** | 0.8204 |
| | | Op. | 54.4885 (127.0829) | 36.1613 (115.5112) | ≪ **0.001** | ≪ **0.001** |
| | $di_{peak}$ | N.Op. | -0.0004 (0.0009) | -0.0004 (0.0011) | ≪ **0.001** | ≪ **0.001** |
| | | Op. | 0.0003 (0.0017) | 0.0008 (0.0023) | ≪ **0.001** | ≪ **0.001** |
| | $I_W$ | N.Op. | 0.0235 (0.0514) | 0.0259 (0.0524) | ≪ **0.001** | ≪ **0.001** |
|  |  | Op. | -0.0005 (0.0890) | -0.0141 (0.1125) | 0.8646 | **0.0010** |

## 4. Conclusions

In this work we have presented a novel framework for assessing the impact of pulmonary resection on pulmonary arterial blood flow. We have achieved this by combining a pipeline that transforms 3D arterial surfaces segmented from CT images into patient-specific 1D computational domains. We then make systematic changes to the computational domains to simulate lung resection. We compute blood flow in all 1646 pulmonary arterial networks. These simulations are successful in 1640 cases. The use of a 1D computational model and the structured tree boundary condition result in low-cost model, so a study of this scale is feasible. Further, the model parameters are physiologically meaningful, so we do not need to tune the model parameters to fit measured data. Once obtained, the simulated flow, pressure, and area are interpolated to have a temporal resolution of 1 ms. Wave intensity is computed for all simulations and the four wave components are found. We perform statistical analyses to show that the model is able to qualitatively capture the behaviours observed by Glass et al. [6] in their clinical study. This is good evidence to support the further development and refinement of this modelling framework.

The key limitation of this work is the lack of patient-specificity. We developed a modelling framework with which we are able to provide evidence that the post-operative changes to wave intensity parameters are attributable to the change in arterial geometry induced by surgery. However, the magnitude of these changes was much smaller than those observed by Glass [8, 6]. We used the same pulmonary flow profile for all simulations, regardless of vessel calibre or count. Repeating this study with patient-specific flow profiles to match the patient-specific vascular geometries may go some way to improving the realism of the simulation outcomes. Further, this information is vital in the event that this framework is implemented in a clinical setting.

Some of resections we have simulated are not clinically realistic. Lobectomy is the standard operation for lung cancer [25] making up around around 81% of lung resections in 2008 [26]. However, the majority of the simulations we have presented would be comparable to a sublobar resection [27]. Changes to wave indices in clinically realistic resections, and a comparison between lobectomy and sublobar resection is an area of active research. We have chosen to make the simulated resections as there are more possible combinations of these than there are possible lobectomies, so the statistics that we produce have greater power.

This model neglects respiration which has been shown to effect pulmonary capillary flow [28]. Further, we do not consider how intraoperative positive pressure ventilation may play a role in lung injury and hence wave intensity changes.

The sample size in this study is small. We used readily available pulmonary surfaces in patients free from respiratory disease to expedite the research process, but similar study with a large sample size may be able to enhance the significance of the results presented here and possibly to give greater insights.

## CRediT authorship contribution statement

**JAM**: Conceptualization, Data curation, Formal analysis, Investigation, Methodology, Software, Validation, Visualization, Writing - Original Draft; **NAH**: Funding acquisition, Project administration, Supervision, Writing - Review & Editing.

## Data Availability

Data will be made available on reasonable request.

## Acknowledgements

This research was supported by the UK Engineering and Physical Sciences Research Council (EP/N014642/1, EP/T017899/1). We thank Prof. Ben Shelley (University of Glasgow and Golden Jubilee National Hospital) and Dr. Adam Glass (Queen’s University Belfast) for instigation of this work, their insight into pulmonary resections and wave intensity analysis.

## Ethics statement

In this study, we analysed human data that had already been collected (with relevant ethical approval) and published. No additional data was collected for this study. Ethical approval was neither required nor sought as this is primarily a computational study.

## Appendix A. Further Model Details

### Appendix A.1. Model Equations

Blood pressure, flow, and lumen area in the large arteries are predicted from a non-linear 1D model based on the incompressible Navier-Stokes equations for a Newtonian fluid in a tapering vessel with thin, elastic walls. A full derivation can be found in, e.g., Mackenzie [29]. In essence, the 1D model is derived from Navier-Stokes (NS) equations in cylindrical polar coordinates via a series of rational approximations. We assume that each vessel in the network in which we simulate flow is much longer than the lumen radius, and that the flow is laminar and swirl-free; these assumptions allow us to neglect the radial and angular components in the NS equations. A 1D computational model is significantly less complex than a 3D model, so are more straightforward to implement, and are less computationally expensive to run. This allows us to easily run many hundreds of simulations seen later in this study.

The *x*-momentum equation

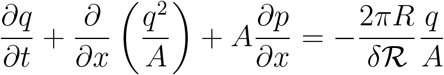

is derived from the NS equations. Here, *t* is time, *x* is longitudinal distance from the inlet of the tube. Reynolds number ℛ and boundary layer thickness *δ* are given. We solve for flow *q*(*x, t*), lumen area *A*(*x, t*), and fluid pressure *p*(*x, t*). Lumen radius is 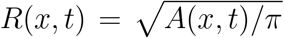. We require two additional equations with unknowns *A, q, p*. These are the continuity equation

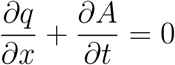

which is derived from the incompressibility condition for elastic walled tubes in cylindrical polar coordinates. Additionally, we also introduce a constitutive relation

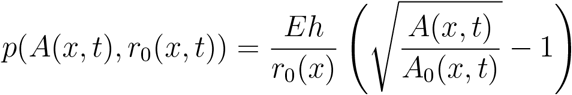

in which *E* is Young’s modulus of the vessel wall, *h* is the wall thickness; *r*_0_ and *A*_0_ are the lumen radius and area in the reference configuration. In practice, these are the radius and area of vessels given when the vessels are specified. Let

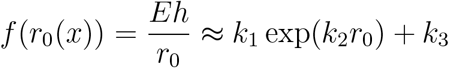

where *k*_1_ = 0, *k*_2_ is arbitrary and *k*_3_ = 26000 Pa for the large pulmonary arteries [24].

The model equations are solved in the network of large vessels using a two-step Lax-Wendroff method. Briefly, the inlet boundary condition (discussed below) is imposed at the proximal end of the MPA which provides *q*(0, *t*) for time *t* over one period. The equations are solved for *A*(0, 0) and *p*(0, 0) given *q*(0, 0). With these, compute *A, q*, and *p* at all spatial steps in the MPA until the distal end is reached. At the distal end, the MPA branches into the left and right pulmonary arteries (LPA and RPA, respectively). The inflow into these comes from the outflow of the MPA. The inflows are computed by imposing continuity of pressure at the boundaries and splitting flow based on the inlet lumen areas of the daughters (see Mackenzie [29]). We now have *q*(0, 0), *A*(0, 0), and *p*(0, 0) for the LPA and RPA; compute *q, A* and *p* at all spatial steps in the interior of these vessels. Once the distal ends of both have been reached, either branch again or terminate with a structured tree as the outlet boundary condition (see main text). More details can be found in, for example, Mackenzie [29].

### Appendix A.2. Structured Trees

In this study, the microvascular is captured using a structured tree approach. This is because the pulmonary arterial system is comprised of many millions of blood vessels that exist on a range of spatial scales, so it is unfeasible so explicitly specify all of these, even if their dimensions and position were known. As it is, imaging modalities have finite resolution that is larger than that of an arteriole [30, 31]. Hence, it is necessary to distinguish between vessels that can be captured in *in vivo* imaging and those that cannot at some spatial scale. However, we cannot simply omit vascular beds from mathematical models, as these are the main site of peripheral resistance. A vascular bed is generated at the distal end of each large vessel that does not give rise to a further vessel. The distal radius of the large vessel from which a vascular bed arises is the radius of the first vessel in the vascular bed. The generation of the vascular beds is discussed in more detail by Olufsen [10]. Vascular beds are truncated at a given radius of 50 µm [24]. The length to radius ratio governs the length of each vessel in the vascular bed; this is chosen as 15.75 [24]. We do not simulate flow or pressure in the vascular beds directly. The vascular beds provide a resistance to flow in the large vessel network; this can be considered to be the down stream boundary condition for the large vessels. At the outlet of the small vessels of the vascular bed, we set a constant pressure of 5 mmHg, the approximate end diastolic pressure in the right ventricle [32].

## Footnotes

1 The offending surface is the penultimate example in the second row of Fig. 3.

2 The custom codebase is run on a single core of a 2022 M2 MacBook Pro with 8GB of RAM. The code base is primarily written in C++ called with a bash script.

